# CDK5 mediated phosphorylation of cytosolic phospholipase A2 regulates its activity and neuroinflammation in Parkinson’s Disease

**DOI:** 10.1101/2021.11.05.467380

**Authors:** Sangita Paul, Saman Fatihi, Srishti Sharma, Rintu Kutum, Raymond Fields, Harish C Pant, Lipi Thukral, BK Binukumar

**Author notes:** Correspondence to: Dr. Binukumar BK, Senior Scientist, Genomics and Molecular Medicine, CSIR-Institute of Genomics and Integrative Biology (IGIB), Mall Road, New Delhi 110007, India.

## Abstract

Hyperactivation of cyclin-dependent kinase 5 (CDK5) by p25, contributes to neuroinflammation causing neurodegeneration in Parkinson’s Disease (PD) and Alzheimer diseases (AD). However, the mechanism by which CDK5 induces neuroinflammation in the PD brain is largely unexplored. Here, we show that CDK5 phosphorylates cytosolic phospholipase A2 (cPLA2) at Thr-268 and Ser-505 sites lead to its activation and generation of eicosanoid products. Mutational studies using site-directed mutagenesis and molecular simulations show that the architecture of the protein changes upon each single-point mutation. Interestingly, double-mutations also led to severe decline in the activity of cPLA2 and disruption of its translocation to the plasma membrane. Further, the brain lysates of transgenic PD mouse models show hyperactivation of CDK5 resulting in enhanced phosphorylation of Thr-268 and Ser-505 of cPLA2 and its heightened activity confirming the findings observed in the cell culture model of PD. These phosphorylation sites of cPLA2 and CDK5 could be explored as the future therapeutic targets against neuroinflammation in PD. Further, conjoint transcriptomic analysis of the publicly available human PD datasets strengthens the hypothesis that genes of the arachidonic acid, prostaglandin synthesis and inflammatory pathways are significantly upregulated in case of the PD patients as compared to that of healthy controls.

## Introduction

Parkinson’s Disease (PD) is one of the most commonly occurring neurodegenerative movement disorders worldwide (Allnutt et al., 2020). It is characterized by a progressive dopaminergic neuronal loss in the substantia nigra forming Lewy bodies (Dauer and Przedborski, 2003). The neurodegeneration is shown to have a strong link with microglial activation and chronic neuroinflammation in the PD brain (Gao et al., 2002; McGeer et al., 1988). Previous studies have reported the dysregulation of CDK5 and increased ratio of p25/p35 in the animal model of PD as well as brains of PD patients (Alvira et al., 2008; He et al., 2018; Smith et al., 2003) along with enhanced neuroinflammation (Kouli et al., 2020; Surendranathan et al., 2015). Neuroinflammation is also previously considered as a key player in PD pathogenesis (Qian et al., 2010), however the precise mechanism behind the link remains unexplained.

Phospholipase A2 (PLA2) is an enzyme that catalyses the hydrolysis of membrane phospholipids from the sn-2 position generating fatty acids such as arachidonic acid (AA) and lysophospholipids. Both molecules are potent inflammatory mediators resulting in immune response (Tan et al., 2016). PLA2 is classified into three groups, namely, secretory PLA2 (sPLA2), cytosolic PLA2 (cPLA2) and Ca2 + independent PLA2 (iPLA2). The cPLA2 or group IV PLA2 have four paralogues known in mammalian cells cPLA2 -α, -β, -γ, -δ out of which cPLA2 α is the one most ubiquitously expressed (Kudo and Murakami, 2002; Litalien and Beaulieu, 2011). PLA2 is a requisite component in the cascade of events leading to the production of eicosanoids during acute and chronic inflammation. Moreover, the prolonged or unmodulated cPLA2 activation leads to membrane deacylation, excessive generation of eicosanoids, and uncontrolled Ca^2+^ influx. These effects ultimately result in lethal cellular injury (Stephenson et al., 1996). Previous evidence implicates inflammatory and immune mechanisms in the pathogenesis of PD and AD (Rogers et al., 2007). There is suggestive evidence that treatment of PD patients with anti-inflammatory agents may be clinically beneficial (Bartels and Leenders, 2010; Thakur and Nehru, 2015).

Hence, the present study is focussed to understand whether CDK5 phosphorylates cPLA2 and, second to determine if phosphorylation of cPLA2 has any effect on its kinase activity as well as translocation to the plasma membrane. The functional significance of CDK5/p25 on cPLA2 activation in primary astrocytes and glia/neuronal culture exposed to MPP+ was also explored. Our studies demonstrate the relevance of CDK5 mediated phosphorylation of cPLA2 in the brain of transgenic PD mice model where activated glia plays a significant role leading to neuroinflammation. Finally, transcriptome analysis of human PD brain dataset compared to its control was performed to study the expression patterns of the genes involved in arachidonic acid and prostaglandin synthesis, inflammatory pathways.

## Results

### CDK5 phosphorylates at S-505 and T-268 of cPLA2

In order to show if cPLA2 is a substrate for CDK5 phosphorylation, Ser-505, Thr-268, and double mutants were generated. Ser-505 and Thr-268 are replaced by Alanine (A) creating cPLA2 single mutants S505A, T268A, and double mutant (S505A, T268A).These plasmids were used for in vitro transcription/ translation for the pure protein synthesis. The pure proteins were used in the CDK5 kinase assay. The kinase assay data (Figure1 A) indicates CDK5/p25 is significantly more effective in phosphorylating cPLA2 wild type(wt) than CDK5/p35. It must be highlighted that both the single mutant, S505A, T268A shows decreased phosphorylation when compared to cPLA2 wt. While, a double mutant of cPLA2 shows decreased phosphorylation compared to the corresponding single mutations. Moreover, the cPLA2 wt phosphorylation is completely halted in the presence of a CDK5 inhibitor. This assay uncovers that S-505 and T-268 are the potential CDK5 phosphorylation sites on cPLA2 (Figure 1A).

**Figure 1:**
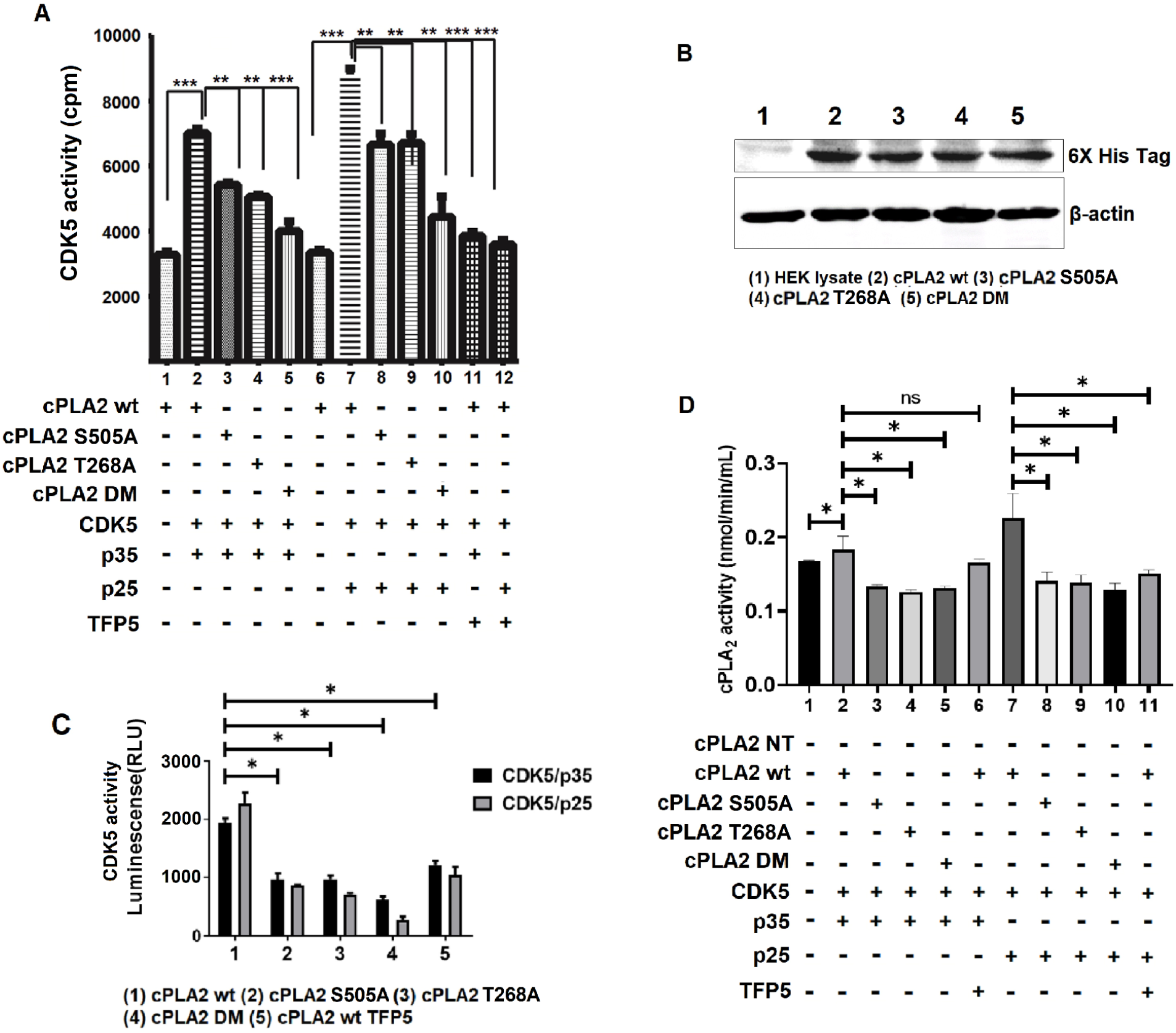
cPLA2 phosphorylation by CDK5. (A) CDK5 kinase assay with cPLA2 wt and its mutants along with TFP5 treatment in vitro by its pure proteins. (**B**) The Western blot image shows the expression of cPLA2 and its mutants post-transfection in HEK293T cells (**C**) cPLA2 wt and mutant protein were used for the CDK5 phosphorylation assay. (**D**). cPLA2 kinase activity: The bar graph shows cPLA2 activity of cPLA2 wt along with its three mutants S505A, T268A and DM, also a comparative analysis of its mutants with the cPLA2 wt activity, ^*^p<0.05

To authenticate above in vitro findings, cPLA2 phosphorylation by CDK5, the expression of the cPLA2 wt, as well as mutants, is confirmed by its transfection in HEK293T cells followed by western blot using 6X-his tag antibody (Figure 1B). The overexpressed cPLA2 wt and mutant proteins were pulled down using immunoprecipitation with 6X-his tag antibody and subjected to CDK5 kinase assay. The in-vitro CDK5 kinase assay results (Figure 1C) also shows decreased phosphorylations in all the cPLA2 mutants. Previously it has been reported that S-505 was a phosphorylating site of MAP kinase. Here, we are first reporting that S-505 and T-268 are the novel CDK5 phosphorylation sites on cPLA2.

### S-505 and T-268 of cPLA2 is critical for its activity

Subsequently, we ensure whether these sites are vital for cPLA2 kinase activity. We overexpressed all the cPLA2 plasmids with CDK5/p35, CDK5/p25 along with the TFP5/SCP treatment. and pulled down the cPLA2 proteins with 6X-His tag for cPLA2 kinase assay (Figure 1C). The result shows a decrease in cPLA2 activity in two single mutants along with the double mutant compared to the cPLA2 wt (Figure 1D). This indicates that both of these sites, S-505 and T-268 are important for the phospholipase activity of cPLA2, therefore mutation in any of the sites or both the sites together affects the cPLA2 activity significantly. Moreover, phosphorylation of these sites enhance the cPLA2 activity (Figure 1D). The CDK5 kinase specific inhibitor treatment shows decreased cPLA2 wt kinase activity.

### Comparison of wild-type and mutant structural cPLA2 using MD simulations

To obtain a deeper understanding of how mutations might affect the functional state of the protein, we performed microsecond-scale molecular dynamics simulations of cPLA2 enzyme. Structurally, the two phosphorylated sites are on different faces of protein, with amino acid position S-505 in the cap region of cPLA2 which is known to be highly flexible(Dessen et al., 1999) and Thr-268 at the nucleophilic elbow. In parallel, we also generated three mutant simulations, namely T268A, S505A, and a double mutant with both positions substituted to an alanine residue. The mutant simulations showed an increase in the angle between two functional domains ∼100°, a significant increase from 120 degrees observed in the wild-type structures. While the non-mutant structures are almost of open configuration (large angular shift), domains in mutant structures are drastically different and represent a more “compact” state.

### Secondary structure evolution of the catalytic domain residues (144-749) according to DSSP analysis in wild and mutant trajectories

In order to further understand local differences, we calculated secondary structural elements and obtained noticeable changes at the active site region (Figure 2C). For instance, the residues between 410-430 occur as coils in wild type which form short helical structures in mutant trajectories. These findings suggested a shift in the contacts due to these critical amino-acid substitutions. Further, we analyzed the network of residue-interaction at each of the mutant sites by quantifying the number of contacts and hydrogen bonds.We observed the number of overall interactions increased for the double mutant system as compared to the wild type. The network topology derived from the structures at 1μs suggests the networks are dense with additional first neighbouring nodes around the central mutant site in case of double mutant. For instance, residue 268 has more neighbouring nodes in the double mutant as compared to the wild type. On a closer look, we found that these results in backbone changes. Rendering the phosphorylated sites according to surface accessibility revealed that the phosphorylated residues which are exposed in the wild type protein are inaccessible in the double mutant protein (Figure 2A&B). Comparison of the phosphorylated Ser-505 residue, which is a major hub in wild type, suggested a loss of interaction with 12 interactions in wild type whereas only 6 were observed in the double mutant. Our network analysis suggests that mutation of these phosphorylating residues results in dynamic changes around the mutant sites which may affect the activity of the protein (Figure 2D&E).

**Figure 2:**
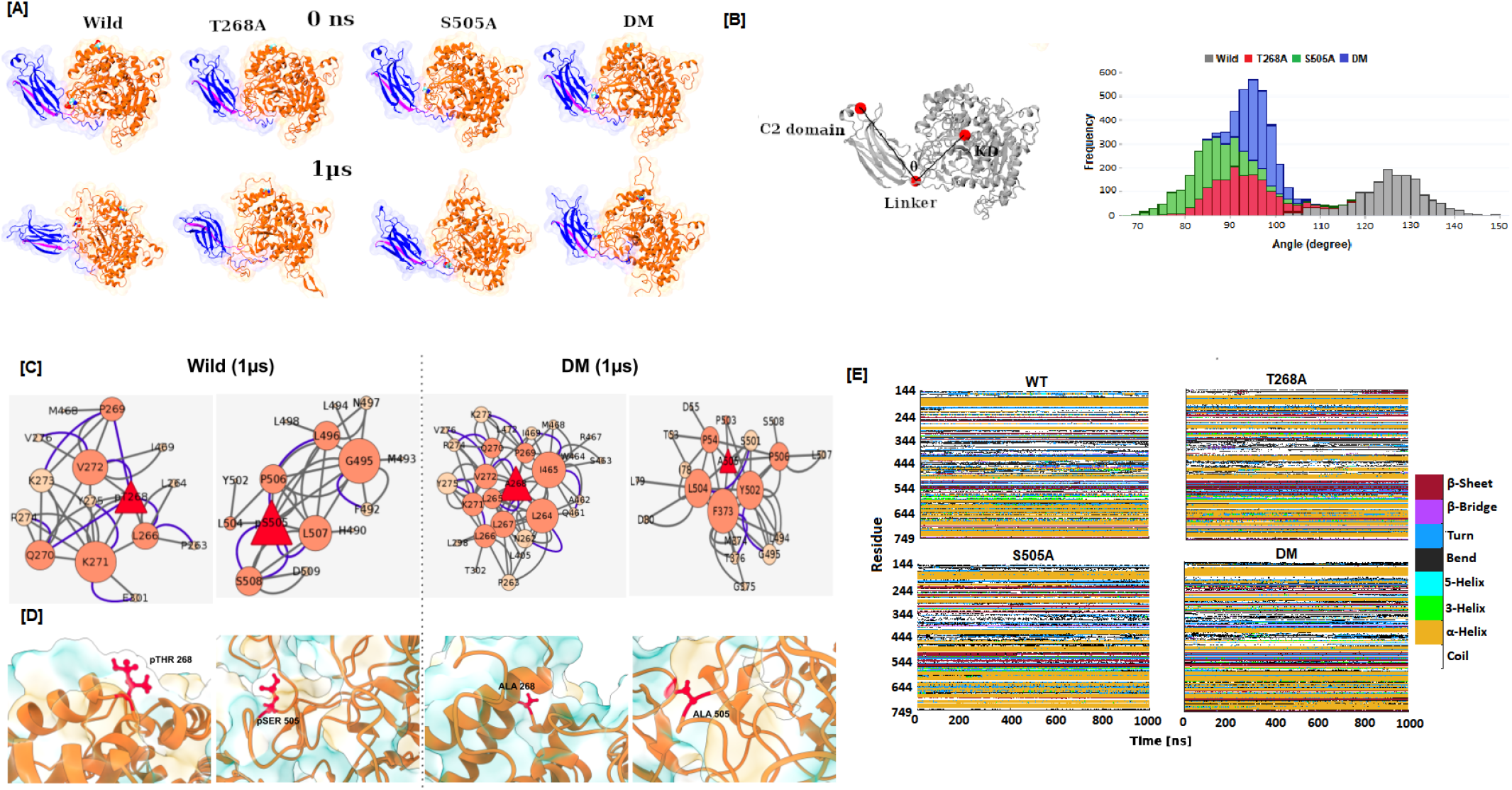
Comparison of wild-type and mutant structural forms using MD simulations. [A] Representative snapshots of MD-simulations derived structure at the initial (0ns) and last (1 μs) of wild and mutant systems. The protein structures are colored according to cPLA2 domain architecture, where C2 domain is highlighted in blue, linker in magenta and catalytic domain (KD) in orange. [B] Angular distribution between domains in wild and mutant systems during the last 150ns of the trajectory. [C] Secondary structure evolution of the catalytic domain residues (144-749) according to DSSP analysis in wild and mutant trajectories [D] Intra-molecular network near the mutant sites for wild and double mutant systems at 1 μs. The size of the nodes represents the degree. The edges are represented as types of interaction, where hydrogen bonds are highlighted in blue and contacts in grey. [E] Snapshots highlighting these sites in ball and stick representation and surface hydrophobicity shown in blue and gold.

### Membrane trafficking of cPLA2 is hindered in its mutants T268A and S505A

Next, we hypothesize phosphorylation on these sites is responsible for the plasma membrane translocation of cPLA2. The hypothesis predicts that cPLA2 is activated upon phosphorylation by CDK5; The membrane and cytosolic protein fractions are collected post-transfection and the expression pattern of cPLA2 wt and mutants are assessed (Figure 3 Using the His-tag antibody for Western analysis of each fraction, the result shows the amount of cPLA2 wt in the membrane fraction is more as compared to cPLA2 T268A, S505A and DM, which are predominantly retained in the cytosolic fraction (Figure 3 A&B). Therefore, cPLA2 phosphorylation at T-268 and S-505 by CDK5 is critical for its translocation from cytosol to the plasma membrane. An antibody for cPLA2 phospho-S-505 is available commercially, however, phospho-T-268 cPLA2 antibody was custom synthesized. Figure 3A shows phospho-T-268 and phospho-S-505 cPLA2 moves to the plasma membrane in the presence of CDK5/p35 and CDK5/p25 at the same time the mutant fails to translocate to the plasma membrane. This is the key experiment which demonstrates that cPLA2 is indeed phosphorylated by CDK5 and phosphorylated cPLA2 at T-268 is translocated to the membrane fraction. In addition, CDK5 inhibitor, TFP5 pretreatment to cells potentiates the effect of CDK5 on the level of plasma-membrane-associated cPLA2 wt (Figure 3D). The increase in the amount of cPLA2 wt associated with the plasma membrane correlates well with the decrease in the amount of cPLA2 wt found in the cytosol following CDK5/p25 and CDK5/35 overexpression, therefore suggesting the translocation of phosphorylated cPLA2 from the cytosol to the plasma membrane upon activation. In the case of cPLA2 S-505 phosphorylation, we also get similar results (Figure 3C).

**Figure 3:**
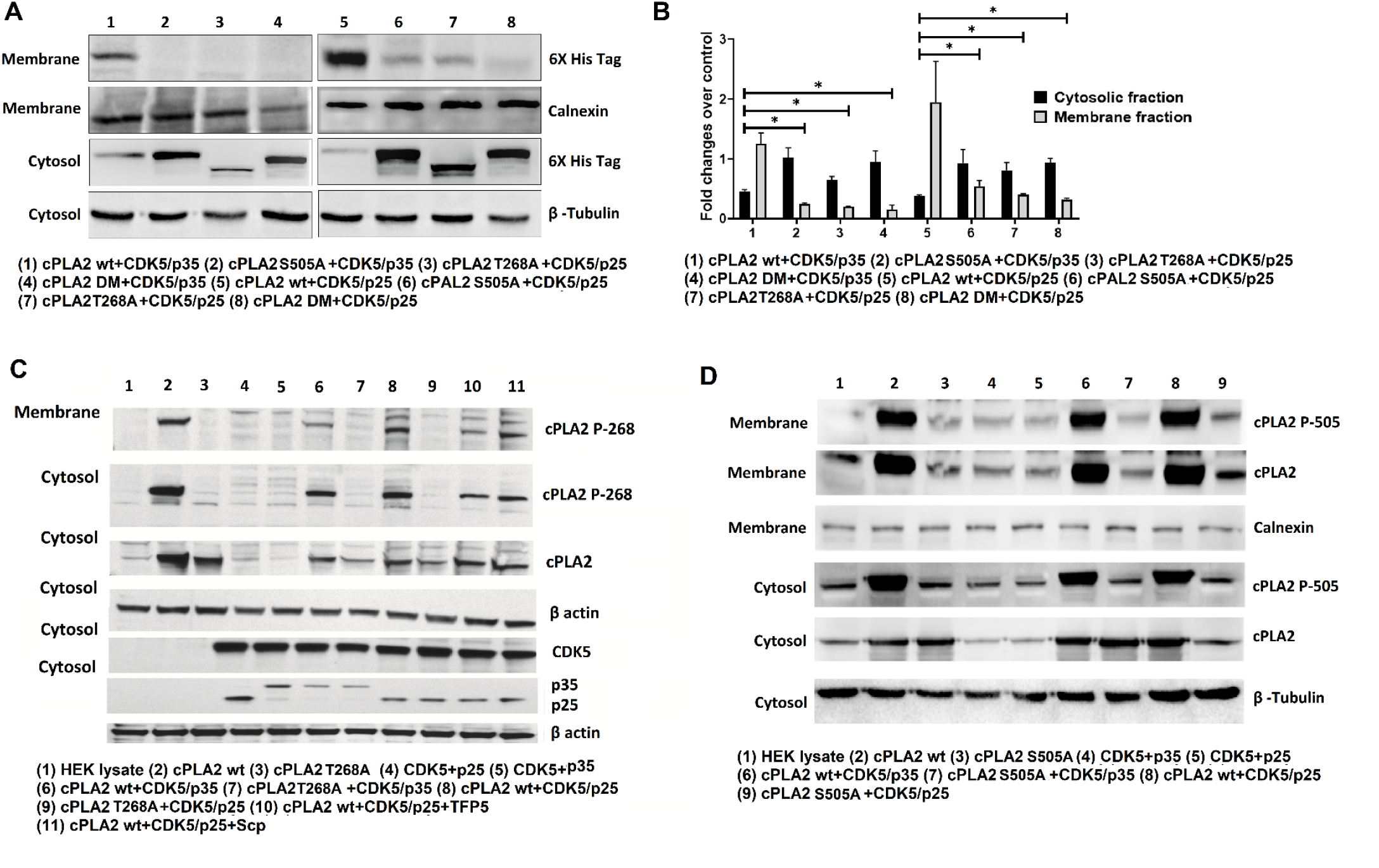
The expression of cPLA2 wt and its mutants in the membrane and cytosolic protein fraction. (A)The expression pattern of cPLA2 and its mutants in a membrane and cytosolic fraction of HEK293T protein lysate co-transfected with cPLA2 and its mutants along with CDK5/p35 and CDK5/p25. The reaction standard used for membrane fraction is calnexin and for cytosolic fraction is beta-tubulin. (B) Cytosolic and membrane fraction fold changes over control. (C) Membrane translocation of phosS505 cPLA2 in the presence of active CDK5 Kinase. (D) Membrane translocation of phosT268 cPLA2 in the presence of active CDK5 Kinase.

### cPLA2 and CDK5 colocalization and interaction

The immunocytochemistry data shows the expression of the cPLA2 wt and mutants in the presence of CDK5/p35 and CDK5/p25 (Supplementary figure 1A). The mutants were predominantly present in the cytosol even in the presence of active CDK5 kinase. Also CDK5 is shown to colocalize with cPLA2. Moreover, co-immunoprecipitation experiments were carried out with mouse brain lysates and CDK5, cPLA2 over expressed in the HEK293T cell lysate. Western blot with CDK5 immunoprecipitation product (IPs) and cPLA2 IPs using antibodies for cPLA2 and CDK5, respectively shows that CDK5 physically interacts with cPLA2 (Supplementary figure 1B)

### Enhanced cPLA2 activity in Primary cortical astrocytes culture exposed to MPP+ via CDK5

Previous studies showed high expression of CDK5 in astrocytes besides its main role in neurons. We also previously reported that MPP+ exposure to midbrain culture leads to the deregulation of CDK5/p25 activity (Binukumar et al., 2014). The primary astrocytes were cultures (Schildge et al., 2013) and pretreated with CDK5 inhibitor, TFP5 (500nM) or scrambled peptide (SCP) for 12 hr. then co-incubated with 10uM MPP+ for 24 hr. We could show that CDK5 is expressed in these cells and the level of expression is unaffected by the treatment nor by the presence of TFP5 (Figure 4 A&B). CDK5 activity, however, was enhanced in the presence of MPP+ and reduced in the presence of TFP5 peptide (Figure 4C). Likewise, p25 generation (Figure 4D) was correlated with increased CDK5 activity (Figure 4C), which was restored to control values after TFP5 treatment. cPLA2 activity was measured after stimulation of primary astrocytes with MPP+ and CDK5 inhibitor pretreatment. We confirmed that cPLA2 was expressed in astrocytes (Figure 4 E&F) and MPP^+^ treatment leads to the two-fold activation of cPLA2 activity compared to untreated cells (Figure 4G). The observed stimulated cPLA2 activity is inhibited by CDK5 inhibitor however SCP pre-treatment has no effect on the stimulated activity of cPLA2 (Figure 4G).

**Figure 4:**
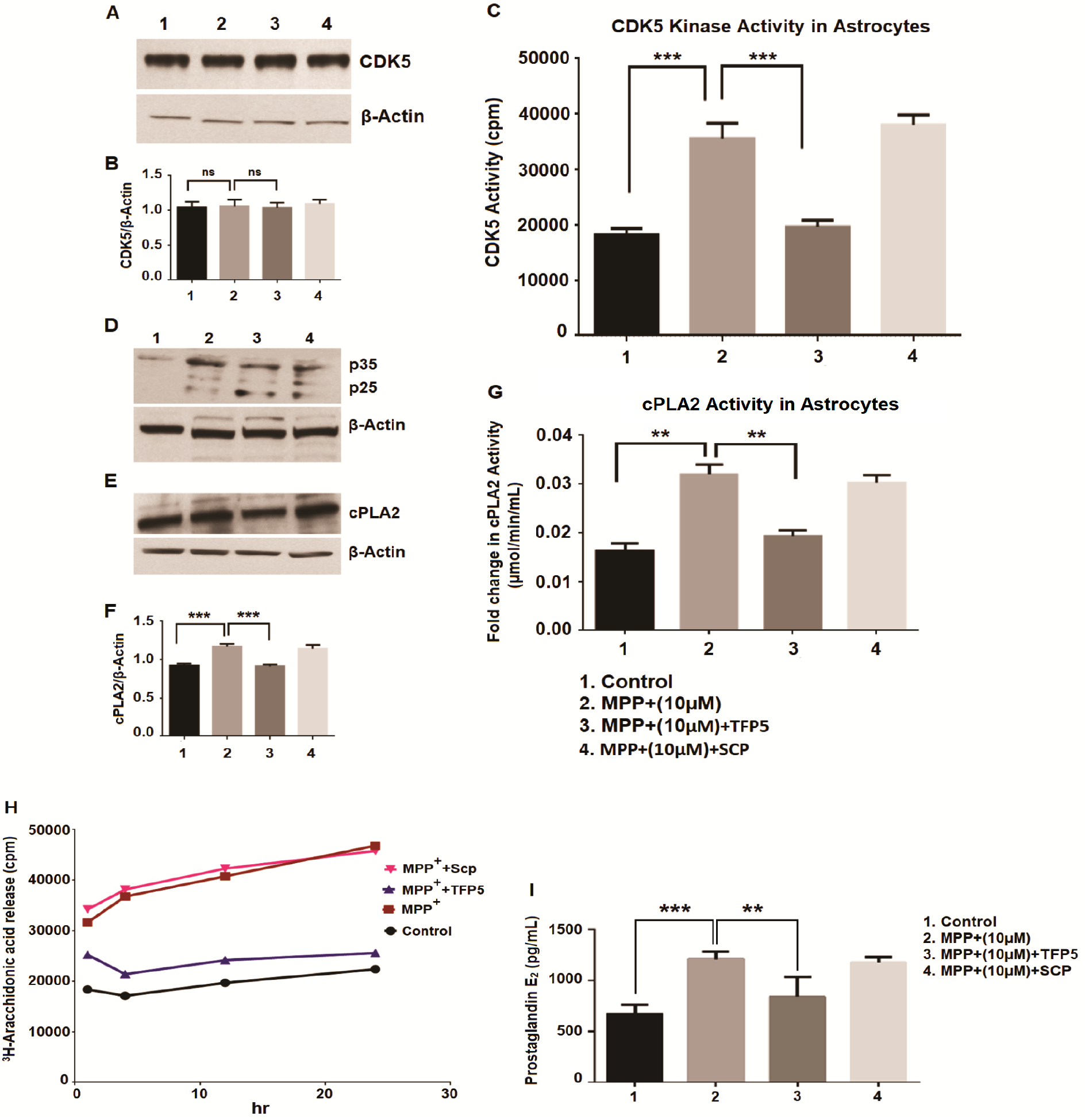
CDK5 expression, generation of p25, its kinase activity, release of arachidonic acid and prostaglandin E2 synthesis by astrocytes exposed with MPP+ and TFP5 treatment. (A) The image shows blot with the expression of CDK5 in control vs MPP+ (10uM) along with MPP+(10uM) vs MPP+(10uM)+TFP5 and SCP treatment in the protein lysate of astrocytes. (B) Bar diagram showing densitometric analysis of CDK5 expression over beta-actin used as a standard for control, MPP+(10uM), MPP+(10uM)+TFP5 and MPP+(10uM)+SCP respectively (C) The bar diagram represents CDK5 activity in the astrocytes protein lysate with control vs MPP+(10uM) and MPP+(10uM) vs MPP+(10uM)+TFP5. (D) The expression pattern of p35 and p25 in control, MPP+(10uM), MPP+(10uM)+TFP5 and MPP+(10uM)+SCP respectively. Beta-actin was used as a standard. (E) The blot representing expression pattern of cPLA2 in control, MPP+(10uM), MPP+(10uM)+TFP5 and MPP+(10uM)+SCP respectively (F) The densitometric graph showing cPLA2 expression over beta-actin of the above blot. (G) The bar graph showing fold changes in cPLA2 activity in astrocytes lysate of control vs MPP+(10uM) and MPP+(10uM) vs MPP+(10uM)+TFP5 (H) The scatter plot showing 3H-Arachidonic acid release from control astrocyte along with MPP+(10uM), MPP+(10uM)+TFP5 and MPP+(10uM)+SCP. (I) The bar graph showing prostaglandin E2 synthesis in control vs MPP+(10uM) and MPP+(10uM) vs MPP+(10uM)+TFP5 along with MPP+(10uM)+SCP as standard for inhibitor treatment. ^*^p>0.05

### Active cPLA2 induces arachidonic acid release and enhanced prostaglandin E_2_ synthesis: initiation of an inflammation cascade is inhibited by CDK5 inhibitor pretreatment

To investigate whether activated cPLA2 releases high levels of arachidonic acid (AA) into the culture medium and TFP5 treatment could reduce its release, we prepared astrocyte cultures from rat brains and quantified AA. Media from cultured cells was collected at different time points (30 minutes, 1 hr, 4 hr, 12 hr and 24 hr) and assayed in a scintillation counter. MPP+ treated cells showed increased AA release compared to control. TFP5 treatment showed significantly less AA radioactivity in the medium compared to the MPP^+^ treatment group (Figure 4H). Figure 4 H&I summarizes the results on the role of potential CDK5 mediated cPLA2 activation and TFP5 in ameliorating factors contributing to neuroinflammation in astrocytes due to MPP^+^ treatment.

To explore prostaglandin E2 (PGE2) synthesis in MPP+ treated cells, the embryonic rat astrocytes were cultured for 12-14 days. The cells were pretreated with/without TFP5 (500nM) or SCP for 12 hr followed by co-incubation in 10uM MPP+, with/wo TFP5 (500nM) or SCP for 24hrs. Prostaglandin synthesis was quantified from the supernatant. MPP+ treated cells showed increased levels of prostaglandin E2 compared to control. CDK5 inhibitor treatment ameliorates this effect and SCP treatment has any effect on the prostaglandin E2 level (Figure 4I)

### Increased cPLA2 activity and enhanced prostaglandin production in neuronal-glial cultures exposed to MPP+ associated with deregulated CDK5 activity

It should be made clear that all experiments described in the pure astrocyte cultures were repeated with mixed neuronal-glial cultures to confirm that the inflammation cascade described is valid for neuron/glia. MPP+ treatment led to enhanced cPLA2 activity in the neuron-glia culture (Supplementary figure 2A&B) and TFP5 treatment showed a decrease in the cPLA2 activity (Supplementary figure 2A&B). Simultaneously, we also measured the PGE2 levels in mixed culture. We notice that similar to primary astrocytes cultures, enhanced PGE2 release in the MPP+ treated cells at the same time TFP5 treatment ameliorates the PGE2 release (Supplementary figure 2C).

### Deregulated CDK5/p25 activity in Transgenic PD mice brain and activation of cPLA2

The expression of p25 and p35 in the brain lysate of the PD mice model was studied with respect to its control along with the CDK5 activity. The result shows enhanced CDK5 expressions along with the generation of p25 in the PD mice brain compared to the control (Figure 5A&B). Also, enhanced CDK5 activity (Figure 5C) supports dysregulation of CDK5 by p25 in the PD brains. Since CDK5/p25 is deregulated, we next checked the cPLA2 activity in the PD mice brain. Figure 5D shows increased cPLA2 activity in the brains of PD mice models compared to control mice. In addition, we could also see the increased level of prostaglandin E2 as well (Figure 5E).

**Figure 5:**
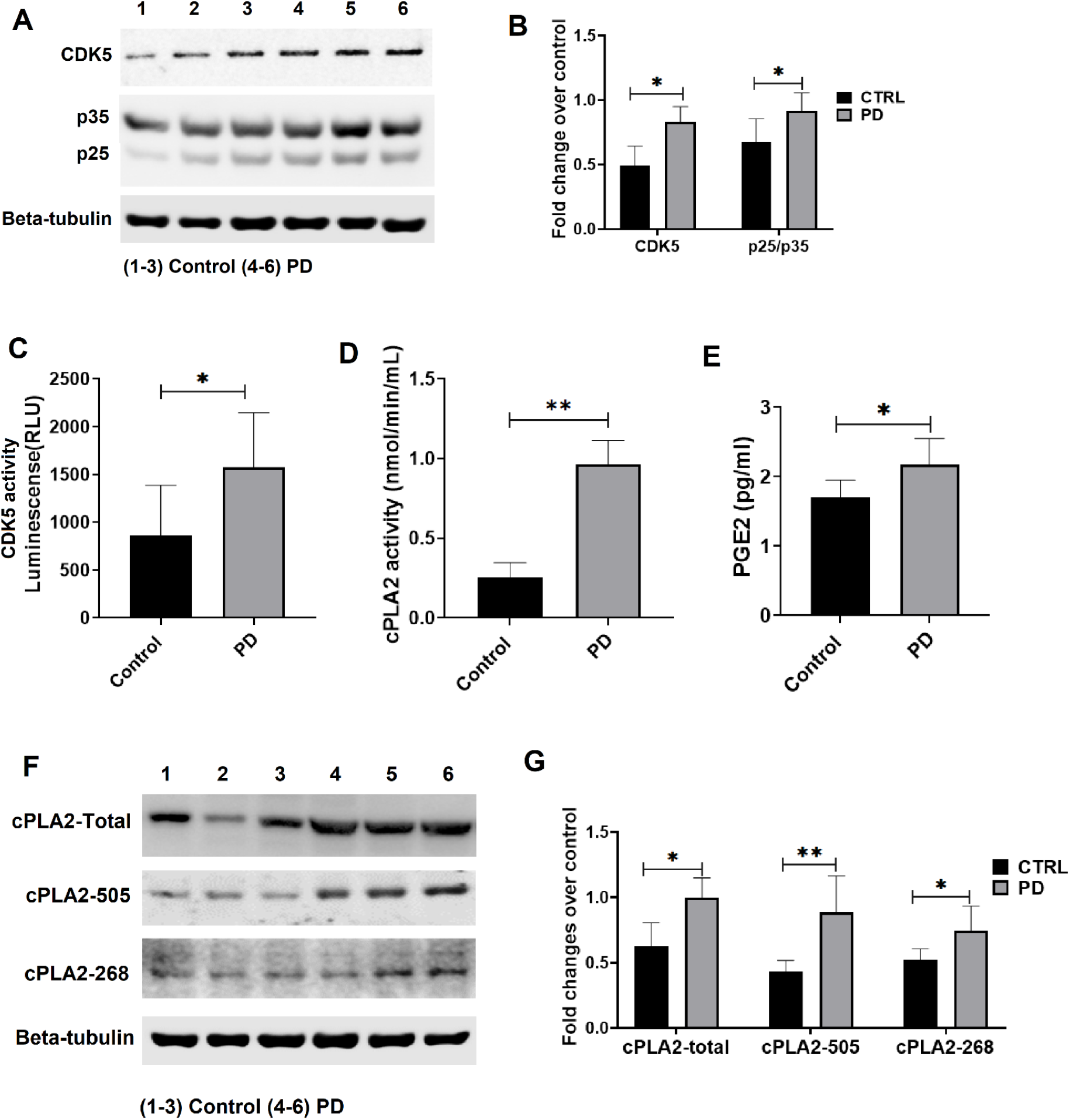
Generation of p25, CDK5 activity, cPLA2 activity and PGE2 levels along with expression pattern of cPLA2 and phospho-cPLA2 in Control vs PD mice model. (A)The blot shows the expression pattern of CDK5, p35 and p25 in Control vs PD mice brain protein lysate (n=3). (B) The densitometric analysis of CDK5 expression and p25/p35 ratio over its standard, beta-tubulin, in control vs PD mice (n=6). (C) The bar graph shows CDK5 activity which correlates with the luminescence(RLU) measure of control vs PD mice brain protein lysate (n=6) (D) The bar graph shows cPLA2 activity in the brain protein lysate of Control vs PD mice model (n=6). (E) The bar graph shows the PGE2 level in pg/ml of control vs PD mice model (F)Western blot analysis of cPLA2-total, cPLA2-505 and cPLA2-268 expression in Control vs PD mice (G) The bar graph shows fold changes of cPLA2-total, cPLA2-505 and cPLA2-268 expression over beta-actin in control vs PD mice. *p<0.05.

### Enhanced expression of cPLA2, Ser-505 and Thr-268 phosphorylation in PD mice brain

Since we found increased cPLA2 activity in transgenic PD mice brain, we further checked the specific T-268 and S-505 phosphorylation levels. The custom made phospho-specific antibody for the site T-268 and commercial phospho-S505 antibodies are used. The brain protein lysate of PD mice showed increased P-505 and P-268 as compared to the respective control mice (Figure 5F&G). In addition, we could see the astrocytes activation in the PD brain (Figure 6A&B). Immunohistochemistry studies also showed the increased expression of cPLA2 and its phospho-forms S505 and T268 in the PD mice brain compared to respective control (Figure 6C-F).

**Figure 6:**
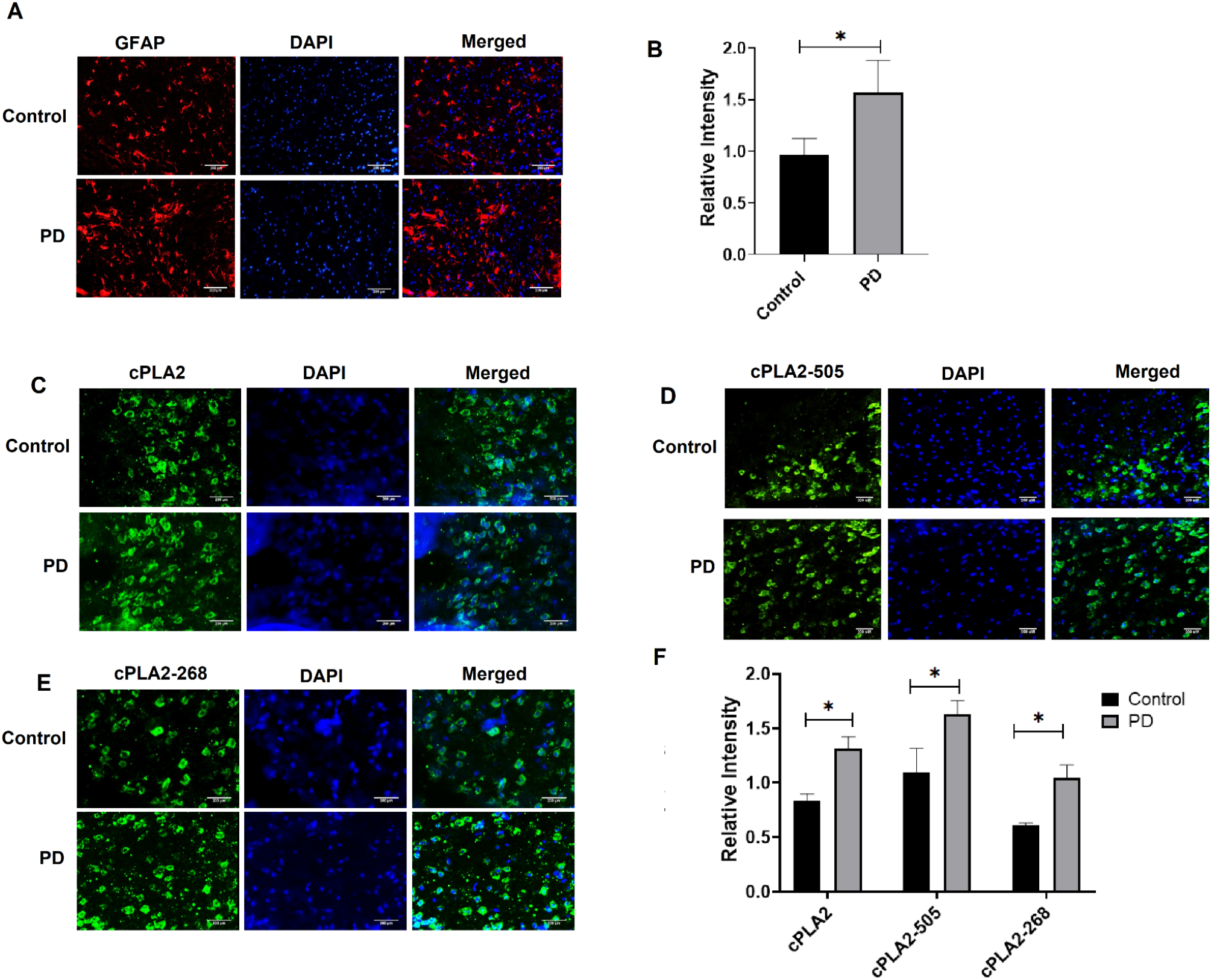
Astrocyte activation along with whole and phospho-cPLA2 (S505 and T268) enhanced expression in the PD mouse brain. (A) Brain sections (10 μm) of control and PD mice were stained with GFAP in control vs PD mice brain section. (B) GFAP immunodensity/mm2 in control vs PD (n =3 mice/group) (C)Expression of GFAP in control vs PD mice brain section (D) Expression of phospho cPLA2 at S-505 (E) Expression of phospho cPLA2 at T-268 in control vs PD mice brain section. Scale bar: 200 μm. (E) cPLA2 and phospho-cPLA2 immunodensity/mm2 in control vs PD(n =3 mice/group), ANOVA with Tukey’s post-hoc test. All data are mean ± SEM. Astrocytes activation and enhanced cPLA2 phosphorylation, S505 and T268 were observed in the PD mouse brain as compared to respective control.

### Human transcriptome analyses of Parkinson’s Disease datasets

To understand if neuroinflammation and other pathways such as prostaglandin and arachidonic acid synthesis are getting modulated in patients with PD, we performed the differential expression analysis of four independent datasets (107 samples) corresponding to substantia nigra of the postmortem samples of the PD(65) and healthy control(42) samples. We queried and performed further analysis with the list of genes of our pathways of interest (AA, Prostaglandin and Inflammatory pathways). Out of the list of 198 genes of the pathway of interest, 67 genes were found to be differentially expressed. Out of this set of genes, 40 genes corresponding to the AA and prostaglandin synthesis metabolism in humans were significantly upregulated whereas 27 genes were seen to be downregulated (Figure 7B and 7C respectively). The genes upregulated included various cytokines and inflammatory pathway genes including interleukins. Also, the various isoforms of phospholipases were seen to be upregulated.

**Figure 7:**
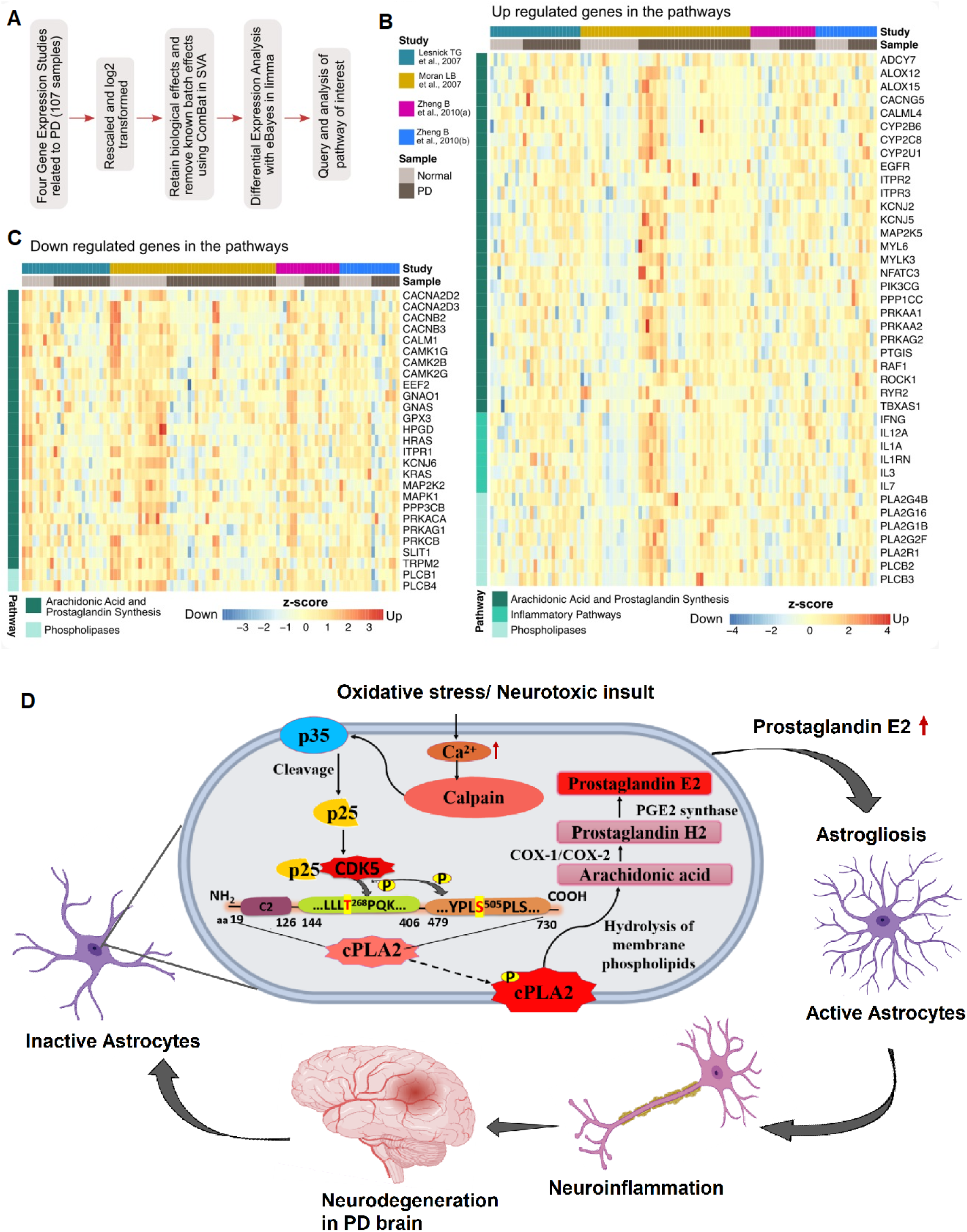
A) Workflow of the conjoint differential expression analysis of four PD transcriptomics datasets. B) Heatmap showing up-regulated genes in the Arachidonic acid and prostaglandin synthesis and Inflammatory pathways. C) Heatmap showing down-regulated genes in the AA and prostaglandin synthesis and Inflammatory pathways. D) The model proposed to explain the mechanism of CDK5/p25 mediated cPLA2 activation leads to neuroinflammation

## Discussion

This study discloses new insights in the role of CDK5 in the regulation of cPLA2 phosphorylation, activity, membrane transport and finally its implication in the PD. The individual single mutant of cPLA2 along with the double mutant fails to phosphorylate in the presence of active CDK5/p35 and CDK5/p25. However, the double mutant does not show an additively decreased level of phosphorylation. Again, in the presence of CDK5 inhibitor TFP5, CDK5 fails to phosphorylate cPLA2 wt (Figure 1A). This confirms that S-505 and T-268 are the potential CDK5 phosphorylation residues on cPLA2. We also checked whether these sites are important for cPLA2 kinase activity. These results are in agreement with the previous data, phosphorylation-dependent increase in cPLA2 activity was observed in cell lysates derived from ATP-treated CHO cells or EGF treated kidney mesangial cells (Bonventre et al., 1990; Goldberg et al., 1990; Gronich et al., 1988; H. Y. Lin et al., 1992; L. L. Lin et al., 1992). Moreover, it has been previously reported that MAPK can phosphorylate S-505 in cPLA2 and a mutation at the consensus site for MAPK in cPLA2 (S505A) fails to be phosphorylated or to generate the decreased electrophoretic mobility form of cPLA2 after incubation with MAPK in vitro (Lin et al., 1993). Through multiple ways, cPLA2 can be phosphorylated and its activation has been regulated. Among these, intracellular Ca2+ have been shown to play an important role in the regulation of cPLA2. Multiple kinases including phorbol ester-activated protein kinase C (PKC), growth factor-activated receptor tyrosine kinases (Bonventre et al., 1990; Goldberg et al., 1990; Gronich et al., 1988; H. Y. Lin et al., 1992; L. L. Lin et al., 1992) and MNK1-related Protein Kinases (Hefner et al., 2000) have shown the phosphorylation and activation of cPLA2. In addition, a number of agents are also associated with increased serine phosphorylation of cPLA2 (H. Y. Lin et al., 1992). No data regarding the threonine phosphorylation of cPLA2 have been previously reported. We report here for the first time that CDK5 phosphorylates T-268 position of cPLA2 in addition to S-505. Moreover, studies with cPLA2 lacking the consensus phosphorylation site for CDK5, Ser-505, Thr-268 indicate that CDK5-mediated cPLA2 phosphorylation is essential for the kinase activity. Our MD studies also revealed large changes in the local structures of the mutant cPLA2 protein (Figure 2). Moreover, structural distortions that we observed in our studies affect the stability of the global structure (Figure 2A). Also changes in the microenvironment of the mutations that we could detect show an important role in protein destabilization (Figure 2B). Our longer-timescale simulations have shown more significant structural changes which could lead to the destabilization of the protein and affect its kinetic activity (Figure 2C&D).

The membrane translocation of phosphorylated cPLA2 at Ser-505 and Thr-268 in presence of CDK5/p35 and CDK5/p25 was also shown, which was significantly decreased in case of single as well as double mutant cPLA2 (Figure 3). In the present study, we also reported MPP+ stimulates CDK5/p25 activation, enhances cPLA2 activity, labelled AA release and enhances prostaglandin E2 production from primary astrocytes culture along with neuroglia mixed culture (Figure 4, Supplementary figure 2). This is in agreement with previous studies that different agents ATP, PMA, or A23187 could evoke the release of labelled AA from cells (Xue et al., 1999). In murine astrocytes AA releases mainly due to cPLA2 activation as Kennedy et al. 1995 reported that C57BL/6J mice are known to have a natural mutation causing a frameshift disruption in the Group IIA sPLA2 gene (Kennedy et al., 1995). Our results agree with the notion that multiple phosphorylation sites are present in cPLA2 (Carvalho et al., 1996; Gijón et al., 1999) and that different protein kinases may regulate its activity including CDK5 (Börsch-Haubold et al., 1998; Geijsen et al., 2000; Gijón et al., 2000).

Phosphorylation of cPLA2 at S-505 by ERK1/2 and/or p38 MAPK has been shown to cause a shift in the electrophoretic mobility of cPLA2 (Gijón et al., 2000; Kramer et al., 1996; Lin et al., 1993) as well as an increase in enzyme activity. It has been also reported that homozygous mice with cPLA2 mutation were significantly resistant to MPTP-induced dopamine depletion as compared with littermate control (Klivenyi et al., 1998). Activation of cPLA2 is observed under pathological conditions where inflammation is present. Since, accumulation and aggregation of α-synuclein (SN) is closely associated with PD pathogenesis (Lee and Trojanowski, 2006), study have also shown that α-SN induces synaptic damage through cPLA2 hyperactivation in α-SN treated primary neurons (Bate and Williams, 2015). A previous study also reports neuroinflammation in the PD rat model through the upregulation of AA signalling mediated by cPLA2 (Lee et al., 2010). A recent study in PD patients reports a significant increase in serum lipoprotein associated PLA2(Lp-PLA2) compared to the control group. This shows a strong association of Lp-PLA2 with the risk of PD pathogenesis, also, Lp-PLA2 serum levels can be used for the detection of PD (Wu et al., 2021). The finding of elevated levels of cPLA2 immunoreactivity in the PD brain supports the hypothesis that there is an active inflammatory process occurring in PD via cPLA2.

However, a previous report suggests that there is a close association between neurodegeneration and p25-mediated neuroinflammation (Muyllaert et al., 2008). Neuroinflammation and neurodegeneration are main pathological features of the PD brain. It was reported previously that inhibition of CDK5/p25 hyperactivation leads to reduced neurodegeneration in the PD mouse model (Binukumar et al., 2015). Lastly we also checked whether CDK5 hyperactivation takes place in the brain of transgenic PD mice. Here, we show the generation of p25 and hyperactivation of CDK5/p25 in the brain of the transgenic PD mice model (Figure 5 A-C). We further checked the cPLA2 kinase activity in the PD mice brain. We observed a significant increase in the cPLA2 activity in PD mice brains compared to control (Figure 5D). Moreover, we also noticed hyperphosphorylation of cPLA2 at S-505 and T-268 residues in the PD mice brain (Figure 5 F&G). We also observed the activation of astrocytes in the tissue sections of the PD mice brain (Figure 6A&B). Finally, activation of cPLA2 leads to the significant increase in the production of PGE2 levels (Figure 5E) in the PD mice brain compared to the control brain.

Our in vitro and in vivo results have pointed out that activated phospholipases induce the AA release and enhanced prostaglandin synthesis, this is validated further with the conjoint transcriptomic analysis wherein significant upregulation in the AA, prostaglandin and inflammatory pathways is noticeable in case of PD as compared to healthy controls (Figure 7B). The levels of the genes such as PTGIS (Prostaglandin synthase), PLA2G (phospholipases) and inflammatory cytokines such as IL1A, IL3, IL12 are higher in the PD patients. Upregulation in the oxytocin signalling pathway is also seen to increase the expression of cPLA2 and also increase the calcium ion flux inside the cell (Soloff et al., 2000). Previous studies have shown that activated microglia contributes to the dopaminergic neuron degeneration in substantia nigra of PD patients (Caggiu et al., 2019) (Karaaslan et al., 2021), reports in murine PD models have also shown upregulation in AA signalling, concordant with these findings increased expression of genes encoding pro-inflammatory cytokines and AA signalling and various other key regulators in these pathways has been found in our analyses. Figure7D, shows a proposed model to explains the mechanism of CDK5/p25 mediated cPLA2 activation leading to neuroinflammation. Inhibiting the phosphorylation of S-505 and T-268 of cPLA2 or decreasing the deregulating CDK5 kinase activity in humans could be a potential therapeutic intervention to rescue the excess neuroinflammation in the PD brain.

## RESULTS MATERIALS AND METHODS

### Starting structure

The protein structure of human cPLA2 was taken from the RCSB PDB Database (PDB ID: 1CJY). The missing residues in the crystal structure present in the C2 and the catalytic domain were modelled using I-TASSER(Yang et al., 2015). The point mutations for the residues T268 and S505 were generated with ChimeraX(Goddard et al., 2018).

### Molecular dynamics simulations

We performed all-atom MD simulations using Gromacs 2018(Lemkul, 2019) and the charmm36 force field(Huang et al., 2017). The water molecules were modeled with the TIP3P model.Periodic boundary conditions were used and long-range electrostatic interactions were treated with Particle Mesh Ewald (PME)(Darden et al., 1993) summation using grid spacing of 0.16 nm combined with a fourth order cubic interpolation to deduce the potential in-between grid-points. The real-space cut-off distance and van der Waals cutoff was set to 1.4 nm. The bond lengths were fixed and a time step of 2 fs for numerical integration of the equation of motions was used(Hess et al., 1997). Coordinates were saved every 20 ps. Four independent MD trajectories, each 1 μs long at 310 K were carried out for cPLA2 wild and mutant structures.The four systems are referred to as wt, T268A, S505A, DM (double mutant).The protein was placed in a cubic water box with a minimum of 1.0 nm of solvent on all sides. The systems were subjected to energy minimization using the steepest descent method. The simulations were subjected to a Nose-Hoover T-coupling bath to maintain the exact temperature(Nosé, 1984). The structures were then subjected to Parrinello-Rahman barostat for pressure coupling at 1 bar(Parrinello and Rahman, 1981).

### Analysis of trajectories

The trajectories were analyzed for comparative structural changes using gromacs analysis toolkit. The secondary structural changes were calculated by the DSSP method(Frishman and Argos, 1995). Further, to understand intramolecular interactions, we utilized a network approach to study quantitative changes occurring near the mutant site across trajectories.The nodes represent the residues and the interactions between residues are edges of the network. A cutoff distance of 0.5 nm was used to define a contact.All molecular images were generated using VMD(Humphrey et al., 1996) and ChimeraX(Goddard et al., 2018).

### Cloning of cPLA2 gene

The plasmid pcDNA3.1-CPLA2 which contains the human cPLA2 gene was obtained from Li-Yuan Chen (NIH/CC/CCMD). pcDNA3.1-CPLA2 was digested with EcoRI + EcoRV and a 236bp linker(GAATTCTGGATTGTGCTACCTACGTTGCTGGTCTTTCTGGCTCCACCTGGTATATGTCAACCTTGTATTCTCACCCTGATTTTCCAGAGAAAGGGCCAGAGGAGATTAATGAAGAACTAATGAAAAATGTTAGCCACAATCCCCTTTTACTTCTCGCACCACAGAAAGTTAAAAGATATGTTGAGTCTTTATGGAAGAAGAAAAGCTCTGGACAACCTGTCACCTTTACTGATATC) oligo containing Ala-268 was ligated into the EcoRI-EcoRV site to create pcDNA3.1-CPLA2A4T. pcDNA3.1-CPLA2 was digested with BmgB1 + PvuII and a 116bp linker-oligo (GACGTGCTGGGAAGGTACACAACTTCATGCTGGGCTTGAATCTCAATACATCTTATCCACTGGCACCTTTGAGTGACTTTGCCACACAGGACTCCTTTGATGATGATGAACTGGATGCAGCTG) containing Ala-505 was ligated into the BmgB1 + PvuII site to create pcDNA3.1-CPLA2A4S. PCR was performed on pcDNA3.1-CPLA2, pcDNA3.1-CPLA2 A4T and pcDNA3.1-CPLA2 A4S to amplify the cPLA2 genes which were ligated into pFB6XHis (made in this lab) to create pFB6XHIS-CPLA2A4T, pFB6XHIS-CPLA2A4T and pFB6XHIS-CPLA2A4S. These plasmids contain hcPLA2 wt, Ala-268 and Ala-505 respectively preceded by 6 Histidine residues which allow for affinity chromatography purification of the proteins. The Ala-505 sequence was removed from pFB6XHIS-CPLA2A4S by restriction enzyme digestion and used to replace the Ser-505 sequence in pFB6XHIS-CPLA2A4T to create the double mutant, Ala268 and Ala-505, pFB6XHIS-CPLA2DM. The 6XHIS-CPLA2 sequences were PCR amplified out of pFB6XHIS constructs and ligated into pDHFR (New England Biolabs, Ipswich, MA 01938) after removal of the E. Coli DHFR gene to create pDHFR-CPLA2, pDHFR-CPLA2A4T, pDHFR-CPLA2A4S and pDHFR-CPLA2DM. The sequences of all constructs were confirmed by plasmid DNA sequencing.

### In Vitro Protein Expression and Purification

The pDHFR-cPLA2 wt, cPLA2A4T (cPLA2 T268A), cPLA2A4S (cPLA2 S505A) and cPLA2 DM plasmids were used for in vitro transcription/translation using the PURExpress in vitro Protein Synthesis Kit. Proteins (wild type, two single mutants and double mutant) were FPLC purified using an NGC FPLC (Bio-Rad, Hercules, CA) a 5ml HisTrap HP column and His Buffer Kit (GE Healthcare Bio-Science Pittsburg, PA)

### In Vivo Protein Expression

The CMV promoter was inserted upstream of the cPLA2 sequences in the pDHFR constructs which allows the plasmids to express the cPLA2 proteins in mammalian cells

### Primary Cortical Astrocytes and mesencephalic cell culture and treatment

Cultures were prepared from the cortical tissues of embryonic day 18.5, as described previously by Schildge, et al. (Schildge et al., 2013). 12-14 days after the first split, astrocytes were pretreated with/wo TFP5 (500nM) or SCP for 12 hr., then co-incubated in 10 μM/ml concentration of MPP+ and with/woTFP5 (500nM) or SCP for 24hr. The mesencephalic neuron-glia cultures were prepared from C57BL6/J mice using a method reported by Binukumar BK et al (Binukumar et al., 2012). The MPP+ and TFP5 treatment are also the same as reported previously.

### Secondary cell culture

Human Embryonic Kidney 293T (HEK293T) cells were cultured in DMEM media (Thermo Scientific), with 10% fetal bovine serum (FBS, Gibco) and 1X penicillin/streptomycin (Gibco). The cells were tested negative for mycoplasma.

### Expression of cPLA2 mutants

The cPLA2 constructs along with CDK5, p35 and CDK5, p25 were co-transfected using lipofectamine-3000 in HEK293T cells. A total of 1.5ug plasmid was added along with the 1X volume of P3000 and 3X volume of lipofectamine-3000 per well in serum-free media. The cells were harvested 24 hrs post transfection using RIPA buffer and 1X protease inhibitor cocktail (PIC). The concentration of the proteins were checked by BCA protein estimation assay.

### Animal handling

All animal experiments were performed according to approved protocols by the Institutional Animal Care and Use Committee of the CSIR IGIB, India.

### Transgenic PD mice

The PD transgenic mice model, Tg(Th-SNCA^*^A30P^*^A53T)39Eric and respective control mice were procured from CCMB Hyderabad, India.

### Generation of phospho-cPLA2(pcPLA2T268) antibody

The pcPLA2T268 antibody was generated by Genscript Antibody Group

### Western blot analysis

The HEK293T cell protein lysate as well as brain lysates from control (*n* = 6) and PD (*n* = 6) mice were prepared as described previously (Shukla et al., 2017). Polyacrylamide gel running, nitrocellulose membrane transfer and detection were performed as reported previously (Shukla et al., 2017).

### Immunoprecipitation of cPLA2 proteins

The protein lysate post transfection in HEK293T cells is subjected to immunoprecipitation using A/G sepharose beads (Thermo). The beads were bound to anti-6X his tag antibody(2ug/ul) overnight at 4°C and then washed three times using a wash buffer (50mM Tris pH 8.5, 1mM EGTA, 75mM KCl). The 6X his tag bound beads were then incubated with the 170ug protein lysate (HEK293T and PD brain protein lysate) for 5 hrs at room temperature. The beads along with its bound antigen were collected by centrifugation at 13,000 rpm in 4°C for 20 minutes and the supernatant was discarded. The collected beads were washed again 3 times with the wash buffer. The product was eluted using urea elution buffer (7M Urea, 20mM Tris pH7.5 and 100mM NaCl).

### cPLA2 activity assay

The cPLA2 assay of the mutants expressed in HEK293T cells along with the cPLA2 expressed in PD brain was performed using cPLA2 assay kit (ab133090, Abcam). 15ul (30ug) of protein lysate samples were added in 96 well plates. The reaction was initiated using 200ul of Arachidonoyl Thio-PC (substrate solution) and incubated for 60 minutes. 10 µl of DTNB/EGTA was added to each well to stop enzyme catalysis and develop the reaction.The absorbance was measured at 414 nm and the cPLA2 activity was calculated as described in the kit protocol. All the reactions were performed in triplicates.

### CDK5 Kinase Assay

CDK5 kinase assay was performed with the immunoprecipitated product with CDK5 antibody in HEK293T as well as brain protein lysate of PD mice model. The assay was based on the previously published paper (Binukumar bk et al, 2014, 15,17,19) and using ADP-GloTM kinase assay kit (Promega)

### Cytosolic and Membrane protein fractionation

The HEK293T cells were harvested 24 hours post-transfection. The cytosolic and membrane protein fractionation was done using the Mem-PER Plus Membrane Protein Extraction kit (ThermoFisher Scientific). The cells from each well were collected using a scraper followed by centrifugation at 300 X g for 5 minutes. The cell pellets were washed with 200ul of wash buffer each twice. The cell pellets were then treated with a 100ul/well permeabilization buffer for 10 minutes at 4°C with constant mixing. The permeabilized cells were centrifuged at 16,000 X g for 15 minutes at 4°C. The supernatants containing the cytosolic fraction were collected. The pellet is further treated with 100ul of solubilization buffer and incubated for 30 minutes at 4°C with constant mixing. The solubilized membrane fraction was then centrifuged at 16,000 X g for 15 minutes at 4°C. The supernatant containing solubilized membrane associated protein was collected.

### PGE2 ELISA assay

The PGE2 elisa assay was done using Prostaglandin E2 ELISA kit (thermo). The reaction mixture for the sample was made in PGE2 coated 8 well strips where 60ug of brain protein lysate samples were added along with 50ul of PGe2-AP tracer and 50ul of PG2 monoclonal antibody supplied within the kit. Along with the sample reaction a standard reaction was also set up for PGE2 with dilutions as required. After 2 hours of incubation 200ul of pNPP solution was added and again an incubation of 70 minutes was done in complete darkness. Absorbance at 405 nm was measured and the PGE2 in pg/ml was quantified in the sample using the calculations given in the protocol.

### Immunocytochemistry

HEK293T cells were seeded in poly-D-lysine coated glass coverslips placed in a 12 well plate. After 12 hours of seeding, the cells were co-transfected with cPLA2 variants along with CDK5/p35 and CDK5/p25 respectively. 24 hours post transfection, the cells were fixed with chilled 100% methanol for 5 min. After fixation the cells were incubated with a blocking solution (1% BSA, 22.52mg/ml glycine in PBST) for 30 min. The cells were then incubated with cocktail primary antibodies cPLA2 (1:250; Sigma Aldrich) and CDK5 (1:250; Invitrogen) for 1 hour in a humid chamber. After incubation with primary antibodies the cells were washed with PBS three times and incubated with secondary anti-mouse 488 nm and anti-rabbit 594 nm (1: 500 each; Invitrogen) for 1 hour. The cells were then washed with PBS three times, mounted with DAPI (Invitrogen) and sealed in glass slides.

### Immunohistochemistry protocol

Ten-micrometer cryostat sections of the brain were collected on slides and prepared for immunohistochemistry, which was performed according to standard protocols for single or double immunostaining (Binukumar et al;2015). Primary antibodies were Pcpla2 S505 (1:100: Sigma Aldrich), Pcpla2 T268(1:50; GenScript Biotech, USA Inc), cPLA2 (1:250; Sigma Aldrich), and GFAP (1:300; Santa Cruz Biotechnology) and GFAP (1:1000; Wako Chemicals). Immunostaining was visualized by Texas red fluorescein and FITC secondary antibodies (Vector Laboratories) and was examined by transmitted or confocal microscopy.

### PD Patients Transcriptome Data Analysis

The conjoint analysis of the PD Patients was carried out in order to check the expression profiles of the arachidonic acid, prostaglandin synthesis pathway and inflammatory pathway genes in the context of humans. We used “Parkinson’s disease” as the search term to screen for the desired datasets on NCBI-GEO and the ArrayExpress database. The datasets were chosen such that they belonged to the substantia nigra of the PD patients. Datasets used in the analysis are as follows : GSE8397(Moran et al., 2006), GSE20141(Zheng et al., 2010), GSE7621(Lesnick et al., 2007), GSE20163 (Zheng et al., 2010). Prior to differential expression analysis, we have performed preprocessing, log2 transformation and normalization according to the downloaded expression matrix. Preprocessing involved 1) common probes expression profiles and 2) rescaling of the data; 21941 common probes corresponding to 13193 genes were used for rescaling expression profiles to 1 (minimum value). Then we performed log_2_ transformation on the profiles. Normalization of the data was performed using surrogate variable analysis. We used the ComBat function of the sva package (Leek et al., 2012) to remove the batch or study specific effects and retain biological effects (expression profile related to PD and healthy control). To inspect the normalization of the expression profiles, we performed principal component analysis on the normalized and without normalization expression profiles (Supplementary Figure 3). The normalized expression profile was further used for differential expression using eBayes (Smyth, 2004) (Johnson et al., 2007) method available in the limma (Ritchie et al., 2015) package in R.

## Supporting information

https://docs.google.com/document/d/1FRq9uF6-a8IVeeZgGcl91qOok2ZddImU4mgKXqund4Y/edit

## Author contributions

BK. conceived of the presented idea. S.P., and B.K. carried out the experiments. S.F. and L.T. performed molecular dynamics simulations. S.S. and R.K. performed transcriptome analysis, R.F. generated cPLA2 plasmid constructs. B.K. wrote the manuscript. H.P., B.K., S.P., L.P. and R.F. contributed to the final version of the manuscript. All authors provided critical feedback and helped shape the research, analysis and manuscript

## Acknowledgment

Authors acknowledge funding support from CSIR and SERB India, and a senior research fellowship to Sangita Paul from CSIR and a junior research fellowship to Srishti Sharma from UGC. Also, we would like to acknowledge Mr. Sanjeev Kumar for his guidance in imaging and Ms. Mukta Poojary for her valuable feedback and insights during the course of study.

