## Supplementary material for "CDK5 mediated phosphorylation of cytosolic phospholipase A2 regulates its activity and neuroinflammation in Parkinson’s Disease": https://docs.google.com/document/d/1FRq9uF6-a8IVeeZgGcl91qOok2ZddImU4mgKXqund4Y/edit

### Supplementary Figures

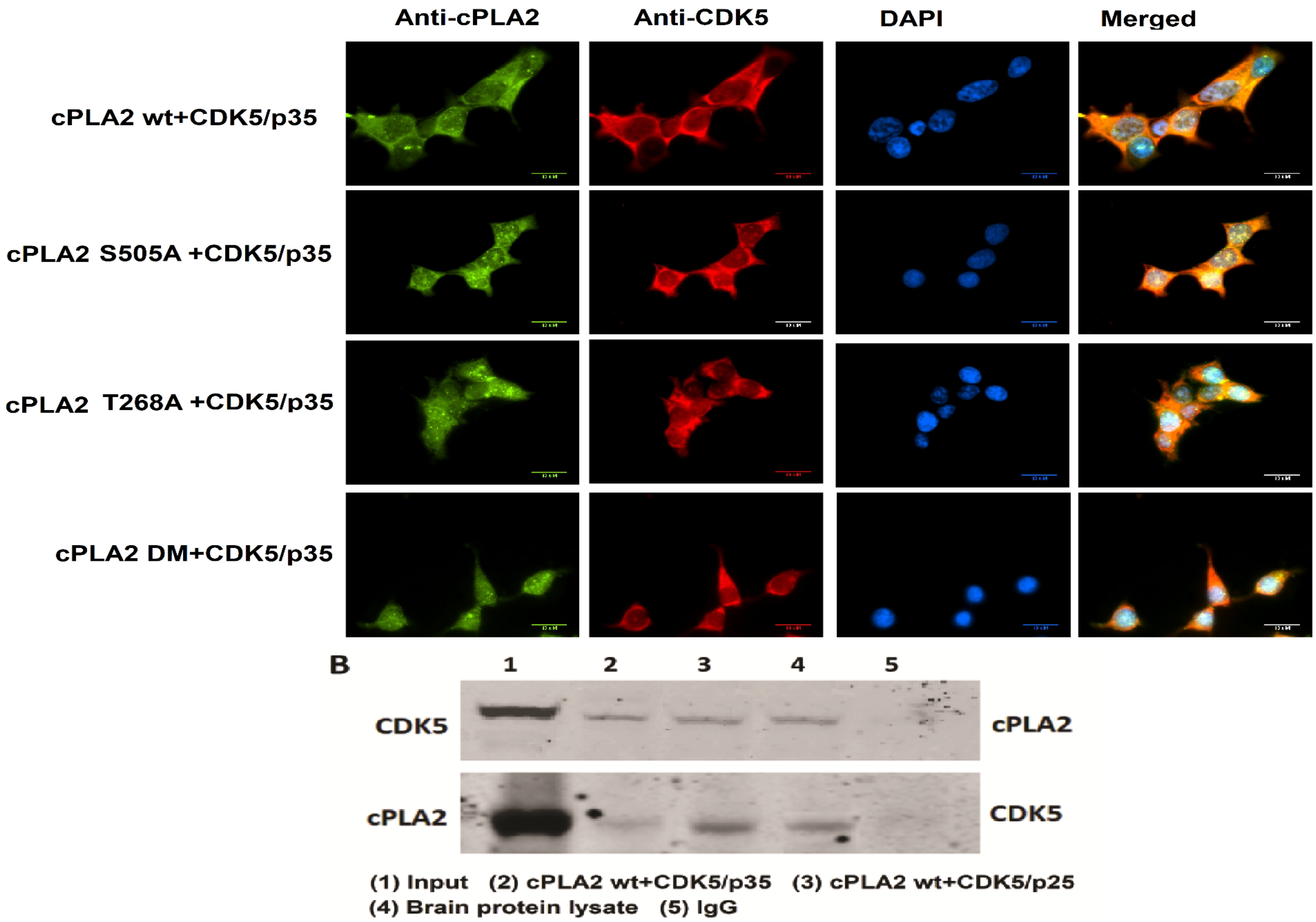


**Supplementary figure 1: Colocalization of CDK5 and cPLA2** (A) Colocalization of CDK5 and cPLA2. Immunocytochemistry analysis of CDK5 and cPLA2 showing their expressions and colocalization. (B) Western blot data of CDK5 and cPLA2 pull down IPs developed with cPLA2 and CDK5 antibodies respectively.

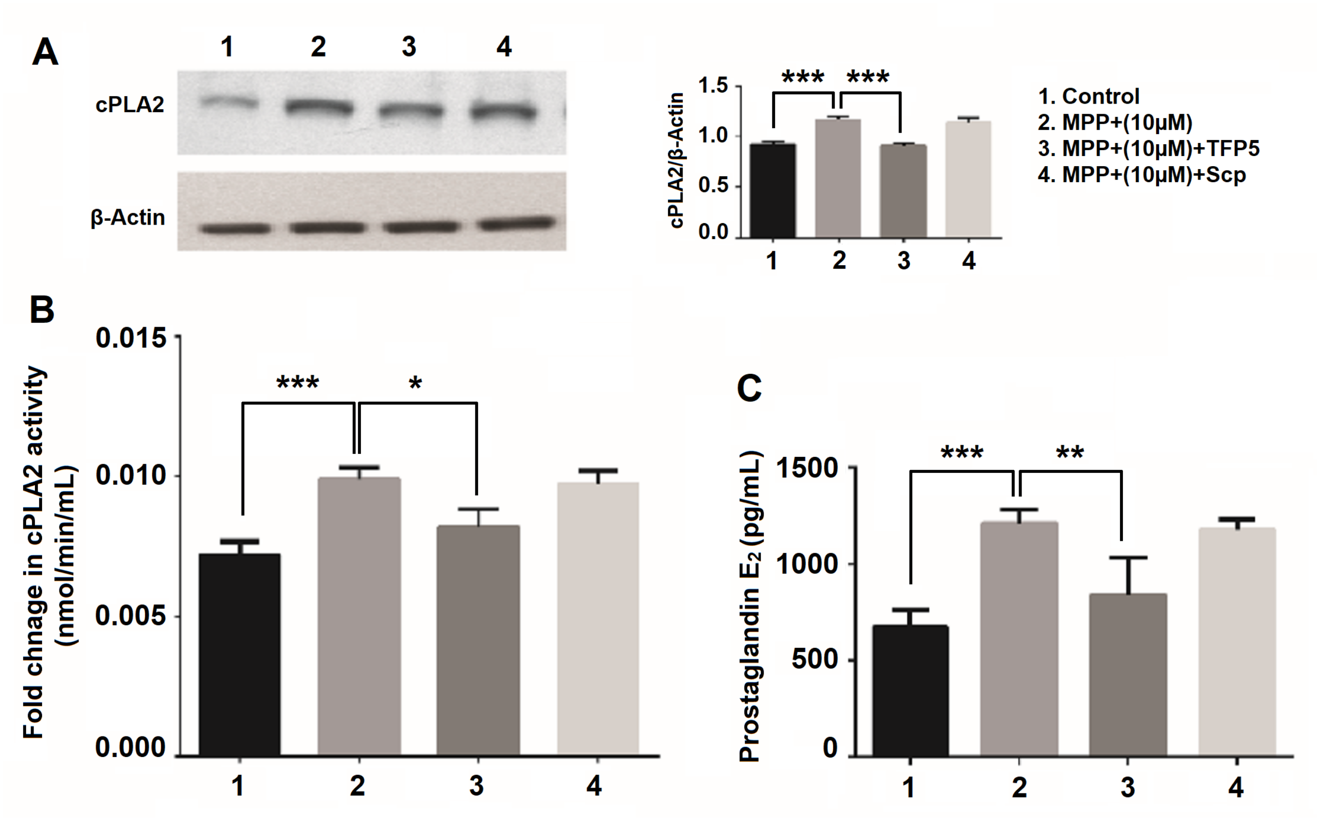


**Supplementary figure 2: cPLA2 expression, its activity and prostaglandin E2 synthesis in neuroglia cells with MPP+ and TFP5 treatment.** (A)The blot showing cPLA2 expression pattern of control, MPP+(10uM), MPP+(10uM)+TFP5 and MPP+(10uM)+SCP in protein lysate of neuronal-glial culture along with its densitometric analysis. (B) The bar graph for the fold of change in cPLA2 activity of control vs MPP+(10uM) and MPP+(10uM) vs MPP+(10uM)+TFP5 in neuronal-glial culture. (C) The bar graph shows prostaglandin E2 amount in control vs MPP+(10uM) and MPP+(10uM) vs MPP+(10uM)+TFP5 in neuronal-glial culture. The MPP+(10uM)+SCP is used as a standard reaction for inhibitor treatment. *p>0.05.

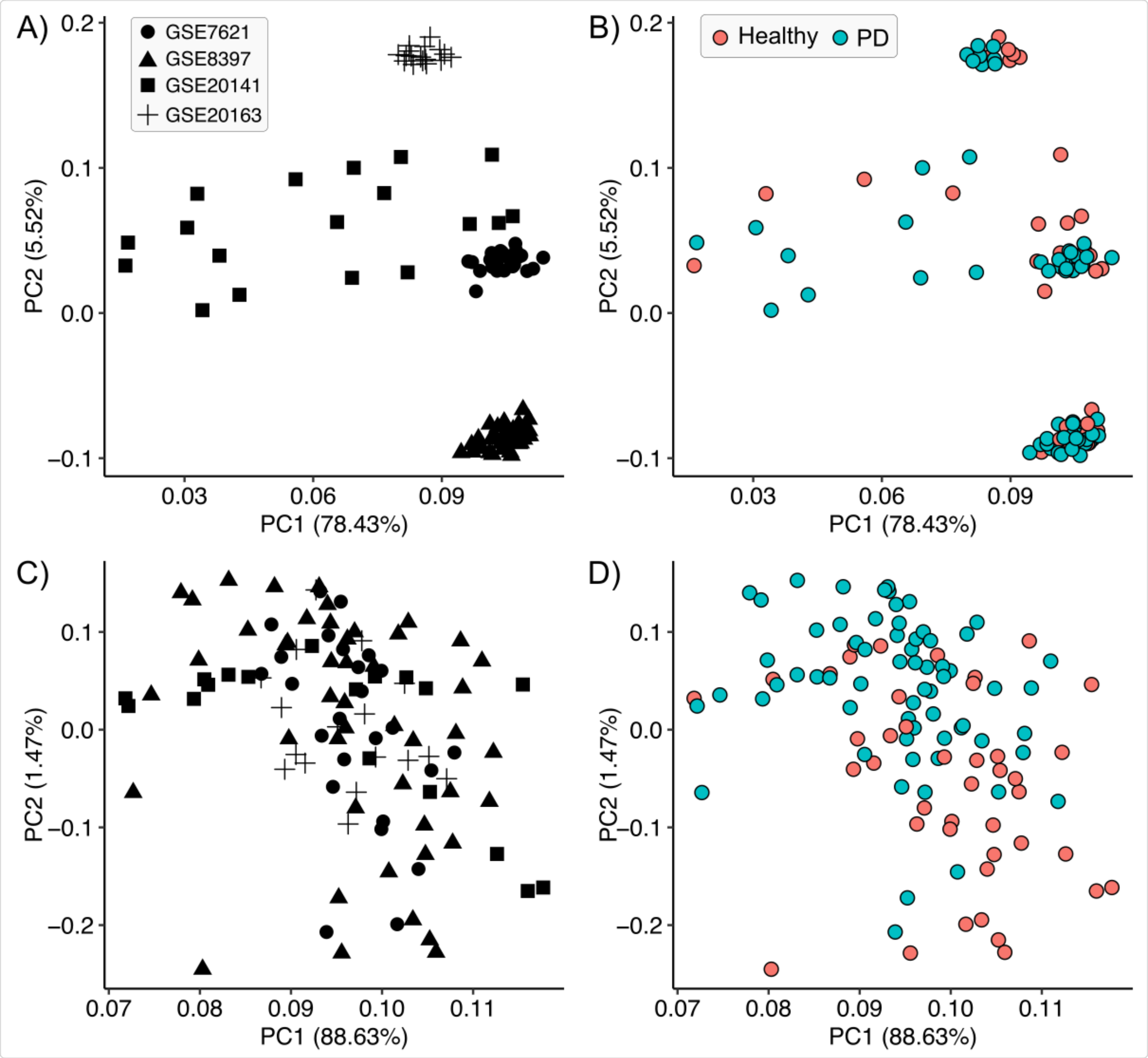


**Supplementary figure 3:** Principal Component Analysis of without (A and B) and with (C and D) normalized expression profiles.
